# Searching for a New Home or Rare Dispersal? Habitat Suitability and Landscape Connectivity of the Tibetan Brown Bear across the Indian Cold Deserts and the Tibetan Plateau

**DOI:** 10.64898/2026.01.21.700975

**Authors:** Vineet Kumar, Amira Sharief, Amar Paul Singh, Shahid Ahmad Dar, Bheem Dutt Joshi, Mukesh Thakur, Lalit Kumar Sharma

## Abstract

Understanding habitat suitability and landscape connectivity is essential for conserving wide-ranging carnivores in climate-sensitive high-altitude mountain ecosystems. The first occurrence record of the Tibetan brown bear (*Ursus arctos pruinosus*) from the Changthang region of Ladakh, India, has raised questions about whether this individual represents an isolated dispersal event or reflects functional connectivity with source populations on the Tibetan Plateau. To evaluate potential habitat suitability and transboundary connectivity, we compiled species occurrence records and associated environmental predictors and developed an ensemble species distribution model using biomod2. We then assessed landscape connectivity using circuit theory implemented in Circuitscape to identify potential ecological corridors. Our models indicate that approximately 1,011,818 km² (21.34%) of the combined Ladakh (India) and the Tibetan Plateau landscape is currently suitable for the species, of which only ∼207,000 km² represents highly suitable habitat. Annual precipitation, precipitation of the wettest month, and precipitation of the warmest quarter were the most influential predictors of habitat suitability. Connectivity analysis identified potential corridors linking the eastern Changthang region of Ladakh with suitable habitat on the Tibetan Plateau, suggesting plausible transboundary ecological connectivity. These results indicate that the recent record from Changthang is more likely driven by landscape-scale functional connectivity than by an isolated dispersal event. Although the mapped corridors represent probable connectivity pathways rather than confirmed movement routes, this study provides the first spatially explicit assessment of habitat suitability and potential transboundary connectivity for the Tibetan brown bear across the Ladakh and Tibetan Plateau landscape.

## Introduction

Climate change, habitat loss and anthropogenic disturbances are the major threats to species persistence. Large carnivores are more sensitive to climate-induced range shifts, contractions, habitat fragmentation and loss due to their extensive spatial requirements and low population densities [1–3]. These factors can have significant negative effects on habitat availability, species distribution and population dynamics [4–8]. Changes in climatic conditions can affect species climatic niche, often evident in geographic range shift and changes in phenological patterns [8–14]. Studies have reported that species respond to adverse effects of climate change mainly by shifting their geographic distributions, and by adjusting through genetic adaption and phenotypic plasticity [15–17]. Shifts in climatic regimes coupled with landscape alteration due to anthropogenic activity can lead to species dispersal into new suitable areas [18, 3, 19). Therefore, identifying and maintaining ecological connectivity across landscapes is essential for facilitating dispersal, population dynamics, and climate-induced range shifts, especially in montane high-altitude ecosystems where suitable habitats are patchy and climate is changing at an unprecedented rate.

Among large carnivores, bears are especially more susceptible to these multi-facet threats. Bears have a broad distribution range, inhabiting four continents, excluding Antarctica, Australia, and Africa [20]. They belong to the order Carnivora and family Ursidae, encompassing eight species globally [21]. These species include the spectacled bear or Andean bear (*Tremarctos ornatus*), Black bear (*Ursus americanus*), Polar bear (*Ursus maritimus*), Brown bear, (*Ursus arctos*), Sun bear (*Helarctos malayanus*), Sloth bear (*Melursus ursinus*) and Giant panda (Ailuropoda melanoleuca) [22,23]. Among all bears species Brown bear (*Ursus arctos*) has large geographic distribution and several sub-species inhabited across North America, Europe, Eastern and Central Asia [22]. Although the Brown bear is globally listed as a Least Concern species by the IUCN, its sub-populations in various regions of Central Asia are considered Threatened [22]. Among the bear species brown bear particularly more vulnerable to the dual threats of climate change and landscape modification due to their large home ranges, long distance natal dispersal and broad habitat requirement. Studies throughout brown bear geographic range has shown adverse effects of climate change on the species [3, 24–33]. In response to climate change and habitat alteration, bears may shift their geographic distribution and expand into new areas, resulting in the detection of species in regions where its presence was previously undocumented. Such observation of Tibetan brown bear has been recorded in camera traps from Changthang Wildlife Sanctuary of Ladakh Union Territory [34].

Earlier in India, Himalayan brown bear is the only sub-species of brown bear known to occupies highlands of the Western Himalayan states of Himachal Pradesh, Uttarakhand, and two Union Territory viz. Jammu and Kashmir Union Territory (JKUT), Ladakh Union Territory (LUT) [35,36]. However, the recent confirmation record of Tibetan brown in Changthang region of Ladakh Union Territory leads to the addition of new bear species to India.

The Tibetan brown bear, also known as the Tibetan blue bear, is a sub-species of brown bears primarily found in the high-altitude regions of the Tibetan Plateau, Nepal, and now also reported from India [34, 37,38]. It is adapted to cold, arid environment of Tibetan Plateau and Trans-Himalaya typically inhabit alpine meadows and scrublands at elevations between 3,000 and 5,500 meters. They are omnivores, and have large home ranges and are known for their ability to travel long distances in search of food. The population on the Tibetan Plateau is estimated to be around 6,300 bears [37]. In this distribution range, Tibetan brown bears frequently come into conflict with nomadic herders and high-altitude communities, as they often attack livestock and break into homes when people are away [39–41]. The similar trends to break into homes was reported in Kungyam village of Changthang region, Ladakh UT [34].

The brown bear population in the Himalayas is mainly distributed in India, Nepal, and Pakistan. The population found in the Upper Mustang Valley [41] and Manaslu Conservation Area in Nepal is thought to be connected to the larger Tibetan population of brown bears and not isolated. However, it is unclear whether the population thriving in the Western Himalayas in the states of Himachal Pradesh, Uttarakhand, and the Union Territories of Jammu and Kashmir, and Ladakh is also connected to the Tibetan population. According to [42] based on genetic studies, there is a historical gap in the range that separates the Himalayan brown bear from the Tibetan brown bear. However, given that brown bears are wide-ranging animals, as evidenced by GPS-collared males and a female with cubs in eastern Tibet, whose home ranges were over 7,000 km² and 2,200 km² respectively [22]. However, the recent camera trap capture of Tibetan brown bear in Changthang region and its proximity to the Tibetan Plateau raises critical and unresolved questions, is it due to climate driven potential range expansion of this species to the Trans-Himalayan region of India, a case of natal dispersal, or is Changthang functionally connected to source populations on the Tibetan Plateau, or does this represent an isolated dispersal event.

For a long ranging species such as bears to move from areas of source population to new suitable unoccupied regions, there is a need of landscape connectivity. Landscape connectivity refers to the extent to which landscape enable or restrict the movement of species between two areas [43]. In the present study we used Species distribution modelling approach widely used to predict potential species distribution and corridor analysis using environmental predictors and species occurrence data to identify suitable habitat, connectivity and range shifts and extinction risks under current and future scenario [44–47]. Further we employed circuit theory-based corridor analysis to identify ecological corridors [26,32,48,49]. In this study, we used occurrence data of Tibetan brown bear from Tibetan Plateau and Changthang region of Ladakh, India, to (i) model suitable habitat for species across a trans-boundary landscape and (ii) applied circuit theory-based connectivity analysis using Circuitscape to identify potential ecological corridor that could connect recently occupied Changthang region with suitable habitat of Tibetan plateau source population.

### Study Area

Our study area included a transboundary high-altitude landscape encompassing Changthang region of Ladakh UT, India and Tibetan plateau (Fig1). This region represents worlds one of the largest continuous high-elevation montane ecosystem, with extreme climatic conditions, low primary productivity, and sparse human settlements [50–53]. Changthang region of Ladakh is form the western extension of Tibetan plateau and share similar ecological and climatic conditions. The regions consist of alpine steppe, cold desert grassland ecosystem, rolling meadows, high-altitude wetlands and rugged mountainous terrain [54]. Climatically, the study landscape comes under rain shadow areas receive low rainfall, high annual precipitation, high temperature variations across seasons, with snow cover persisting for extended periods at higher elevations [55–56]. The vegetation primarily consists of alpine meadows and steppe grasslands, featuring hardy grasses, sedges, and low-lying shrubs adapted to the harsh, cold arid climate [54,57]. The region has the unique assemblage of biodiversity, and home of various rare high-altitude fauna and flora. Major fauna inhabits in the study landscape are, Snow leopard (*Panthera uncia*), Himalayan grey wolf (*Canis lupus laniger*), lynx (*Lynx lynx isabellina*), and Wild dog (*Cuon alpinus laniger*), Blue sheep (*Pseudois nayaur*), Tibetan argali (*Ovis ammon hodgsoni*), Kiang (*Equus hemionus kiang*), Tibetan gazelle (*Procapra picticaudata*), Tibetan antelope (*Pantholops hodgsoni*) and wild yak (*Bos grunniens*), Himalayan marmot (*Marmota himalayana*), woolly hare (*Lepus ostioleous),* Pallas’cats (*Otocolobus manul*)Tibetan snowcock (*Tetraogallus tibetanus*) and chukar (*Alectoris chukar*). The livelihood of the residents of the Changthang region largely depends on livestock rearing, with most local communities engaged in pastoralism, particularly nomadic practices. However, they also do agriculture practice in small scale, even though the region is beyond the usual altitudinal range for farming. Despite its apparent remoteness, the region is experiencing increasing anthropogenic pressures, particularly from pastoralism, infrastructure development, and changing land-use patterns, which may influence habitat availability and landscape permeability for large mammals

**Fig. 1.**
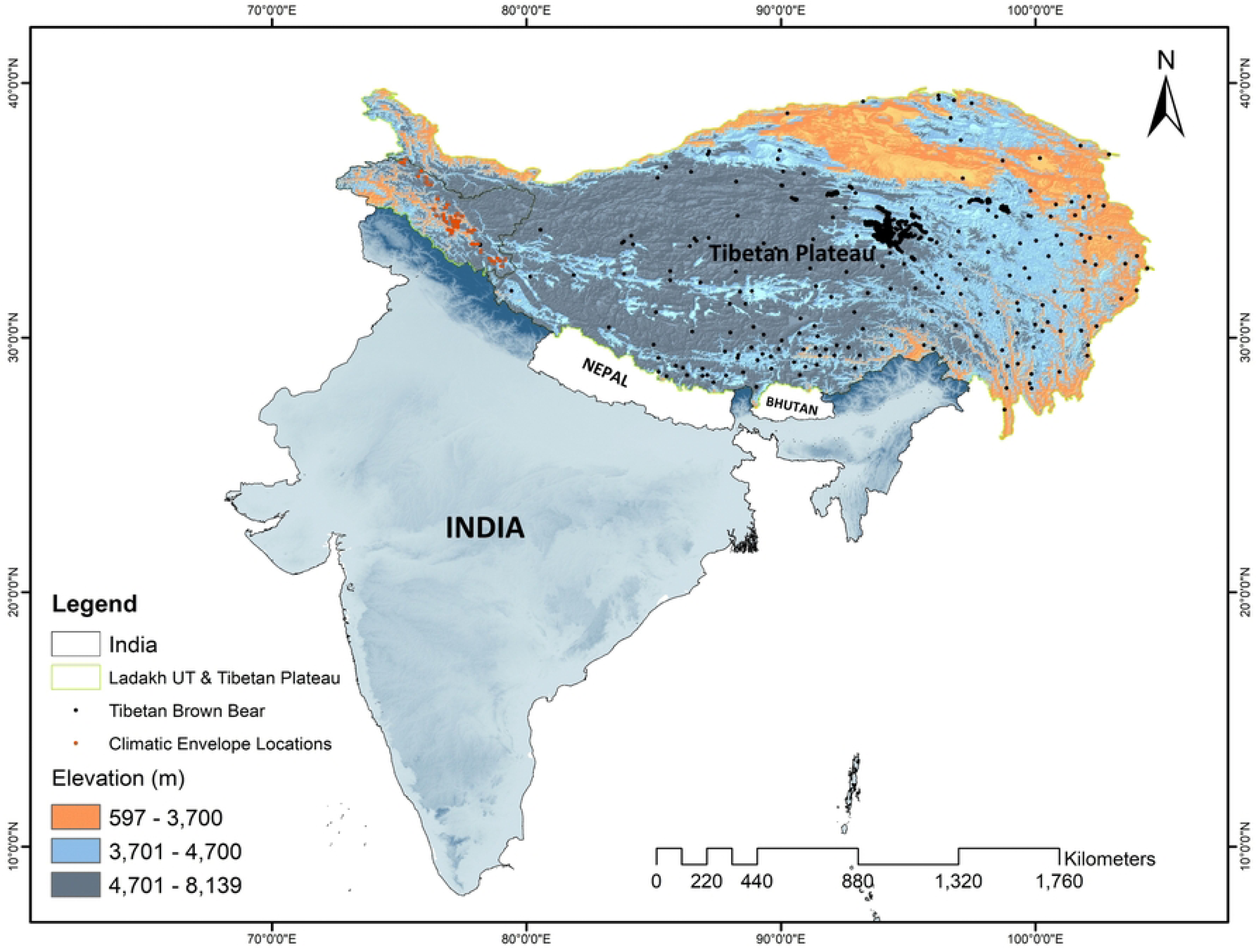
Location of study area (Ladakh UT, India and Tibetan Plateau) with species occurrence record

## Methods

### Presence locations

Occurrence data of Tibetan brown bear were obtained through three camera trap records in Changthang region, Ladakh [34] and 824 published records from the Tibetan plateau [25, 26, 32,37,58, 59]. Tibetan brown bear recorded first time from Changthang region of Ladakh UT with only 3 confirmed locations. Due to low occurrence data on Changthang region, we generated additional 50 pseudo-presence locations only within Ladakh region using an environment envelop sampling approach. Environmental envelope of a species refers to the range of environmental conditions under which it can persist and maintain viable populations, where its ecological requirements are adequately met [60]. In the present paper, we narrow down the concept of environmental envelope to the climatic envelope of a species by identifying the range of climatic conditions within which the species is expected to persist. We hypothesized that potential areas of Tibetan brown bear occurrence will be similar to the climatic conditions where actual presence records of species recorded [61,62]. We used the climatic conditions data of recorded presence locations and created a climatic envelope, then randomly generated the pseudo-presence locations in the areas within climatic envelope where species presence was not observed but can be found [61, 63]. We rarefy the presence records of Tibetan brown bear to 1 occurrence per km^2^ to reduce spatial autocorrelation [64]. The spatial autocorrelation of the locations was tested at 1 km using the SDM toolbox in ArcGIS 10.9. Out of total only 703 spatially independent occurrence records of Tibetan brown bear including climatic envelope locations were used for modelling the distribution in the transboundary landscape (Fig1).

### Environmental variables

In present study initially we used 21 environmental variables (bioclimatic, topographic, and anthropogenic) for modelling the distribution of Tibetan brown bear based on species ecology, geographic features and those used in previous studies [25,26] (S1 Table). 19 bioclimatic variables were extracted from Worldclim Ver. 2 (www.worldclim.org) with ∼1 km resolution [65]. We used SRTM (Shuttle Radar Topographic Mission) image downloaded from Earth Explorer for preparing the topographic variables (elevation) https://earthexplorer.usgs.gov/. Literature suggests brown bear is sensitive to anthropogenic disturbances hence we also used the global Human Influence Index (HII) dataset downloaded from the Socioeconomic Data and Applications Centre SEDAC, NASA (https://sedac.ciesin.columbia.edu). All the variables were resampled with the resolution of 30 arcsec ∼ 1 km spatial resolution using the spatial-analyst tool in ArcGIS 10.9. The Pearson correlation test was performed to identify and remove variables exhibiting significant collinearity. Variables with Pearson coefficient greater than 0.7 (rs>0.7) were dropped from further analysis [66]. Finally, 14 environmental variables which we assumed might have ecological effect on the distribution of the species were retained for modelling habitat suitability of Tibetan brown bear (Table 1). Pearson correlation plot of all the predictors used for modelling (variables with values >7 was excluded from analysis (S1 Fig.)

**Table 1.** Environmental predictors used to model habitat suitability and their contribution.

| Code | Environmental variable | Unit |
| --- | --- | --- |
| Bio2 | Mean diurnal temperature range | °C |
| Bio3 | Temperature isothermality ($\text{Bio2/Bio7} \times 100$) | — |
| Bio6 | Minimum temperature of coldest month | °C |
| Bio9 | Mean temperature of driest quarter | °C |
| Bio11 | Mean temperature of coldest quarter | °C |
| Bio12 | Annual precipitation | mm |
| Bio13 | Precipitation of wettest month | mm |
| Bio14 | Precipitation of driest month | mm |
| Bio15 | Precipitation seasonality (coefficient of variation) | — |
| Bio16 | Precipitation of wettest quarter | mm |
| Bio17 | Precipitation of driest quarter | mm |
| Bio18 | Precipitation of warmest quarter | mm |
| Bio19 | Precipitation of coldest quarter | mm |
| Elev | Elevation | m |

### Species distribution modelling

SDMs are commonly used to model species-environment relationships and forecast species spatial distributions [45,67–70]. Recent advancements, such as the use of computer-based algorithms with varying ensemble procedures produces more reliable habitat suitability maps [71–75]. Ensemble species distribution modelling leverages the strength of multiple models to improve predictive accuracy, handle uncertainty, reduce bias, and provide robust predictions for informing conservation and management decisions. Hence the present study implemented ensemble modelling approach, using R package (‘biomod2’) [76] for modelling distribution of brown bear in the Tibetan landscape. The modelling procedure is summarized into four steps:

1. Formatting the data which includes putting data into the required modelling format and to better simulate the actual distribution and reduce the spatial deviation, 703 pseudoabsence points were randomly selected which was repeated 3 times for model construction [77].
2. Building the model: Three modelling algorithms viz., generalized linear modelling (GLM), (Random Forest (RF), and GAM (Generalized additive model) were used for predicting the distribution of Tibetan brown bear. For selecting best-fitted model with value equal to 0.75 was kept as AUC threshold for finally building the ensemble probability surface. The models were fitted using default biomod2 modelling options. The model calibration was done on 70% of the data and 30% for validation, respectively. We used 10-fold cross validation approach, given that it is equally efficient in the error estimation as other techniques. The model performance was evaluated using the model evaluation score (Receivers Operating Curve (ROC) and True Skill Statistic (TSS). The variable importance scores were extracted from the models using in-built ‘biomod2’ functions. The mean score of variable importance scores obtained with respect to the algorithms were taken into consideration.
3. Building ensemble of models: After generating all model outputs, the best performing models were used to develop the ensemble model, which is the robust strategy to investigate the species distribution [28,78,79]. We verified our final ensemble model by generating ROC (Receiver Operating Characteristic) value, which directed the model evaluation poor when the value is 0.6-0.7, normal with 0.7-0.8, good with 0.8-0.9 value and best with 0.9-1.0 values. Hence, we rejected the models with ROC less than 0.75 value [80].
4. Making projection of models: We produced distribution map of Tibetan brown bear for current scenario in the study landscape. The continuous habitat suitability map was converted into binary probability surfaces with values ranging from 0 ‘absence’ to 1 ‘presence’ by applying a cut-off value which was considered as a threshold. We used biomod2’s built-in binary transformation using binary.meth = "ROC". The threshold (cutoff) was automatically derived by the internal function used by biomod2 (bm_FindOptimStat with metric.eval = "ROC"). Probability surface above the threshold value was classified as suitable habitat otherwise non-suitable.

### Connectivity analysis

We used Circuitscape to assess connectivity between Tibetan plateau and India, leveraging its circuit theory to map ecological connections between habitat patches (Circuitscape software 4.0; https://circuitscape.org/; [26,48,49,81–84]. Circuit model, which is a widely used approach for animal corridor design [85]. The suitable patches were extracted from binary habitat suitability map and then converted to polygons to use as connectivity nodes. Furthermore, the continuous habitat suitability (HS) values were converted into a resistance surface using the negative exponential function R = 1000^(−1 x HS), following the approach of previous connectivity studies [86], where R represents the cost resistance value assigned to each pixel and HS represents habitat suitability. The scaling constant 1000 adequately captures the non-linear biological response of animal movement to declining habitat suitability. This resistance raster reflects the relative cost of movement across the landscape. Output maps depict current flow, where high current flow indicates key areas likely facilitating movement between habitat patches [87].

## Results

### Model Performance

Model selection was based on ROC values greater than 0.75. RF had the highest average ROC value (0.91) amongst all other algorithms (Fig 2). The ROC weighted mean value of the ensemble model was equal to 0.93 which depicts the best performance of all the models used for predicting the distribution of the species.

**Fig 2.**
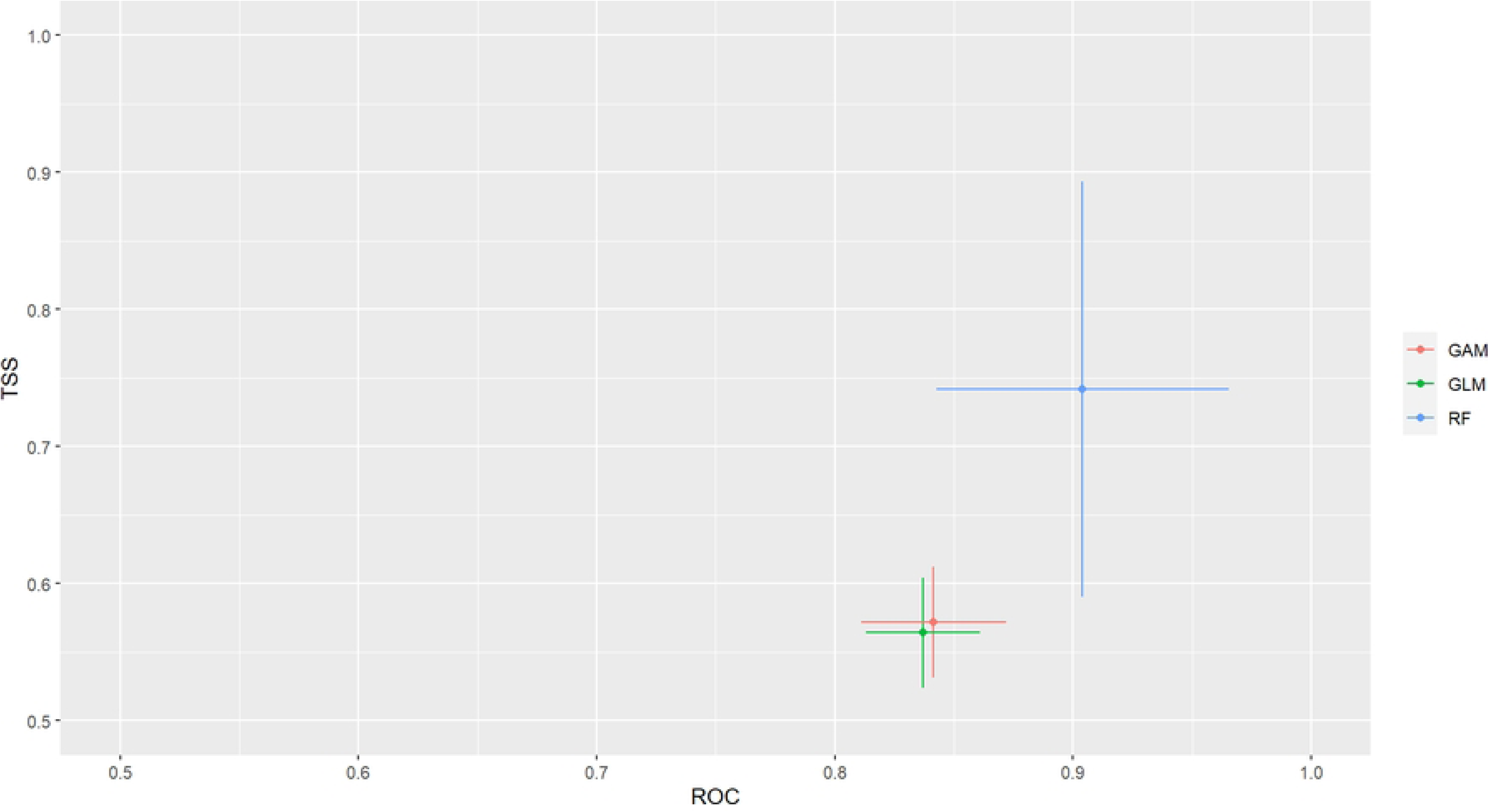
ROC plot of models used for distribution modelling of Tibetan brown bear in the landscape.

Ensemble modelling approach was employed for predicting suitable habitat of Tibetan brown bear in the study landscape. The present study revealed that out of 1011818 km^2^ of the suitable habitat 483487 km^2^ (47.78%) is the low suitable habitat and 321331 km^2^ (31.76%) falls under medium suitable and 207000 km^2^ (20.46%) high suitable area for Tibetan brown bear (Fig 3). Among all the participatory variables, annual precipitation (Bio12), precipitation of wettest month (Bio13) with variable importance equal to 0.48 and 0.41 respectively, were the top influential factors governing the distribution of Tibetan brown bear (Fig 4). The response curve depicts that Bio12 is having positive influence on the distribution, whereas Bio13 showed negative influence on the distribution of Tibetan brown bear. The response curves of all the models used for building ensemble model are presented in below please see (S2 Fig. a-c). Projected corridor for Tibetan brown bear under current climate scenario shows the habitat connectivity from east of Changthang region to the Tibetan plateau (Fig 5). Under the current climate scenario there is high connectivity among the core habitat patches across Tibetan plateau.

**Fig 3.**
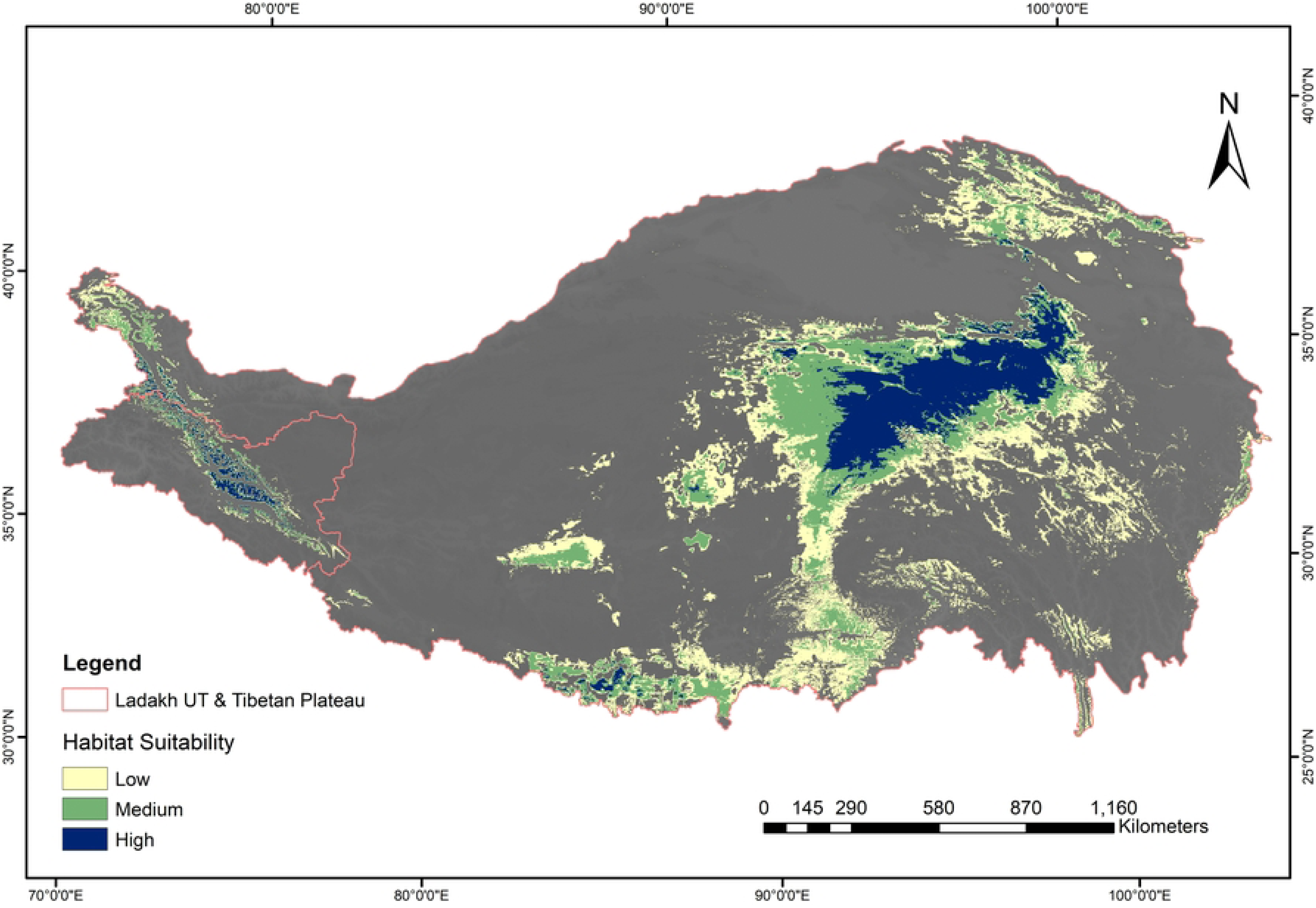
Current habitat suitability map of Tibetan brown bear

**Fig 4.**
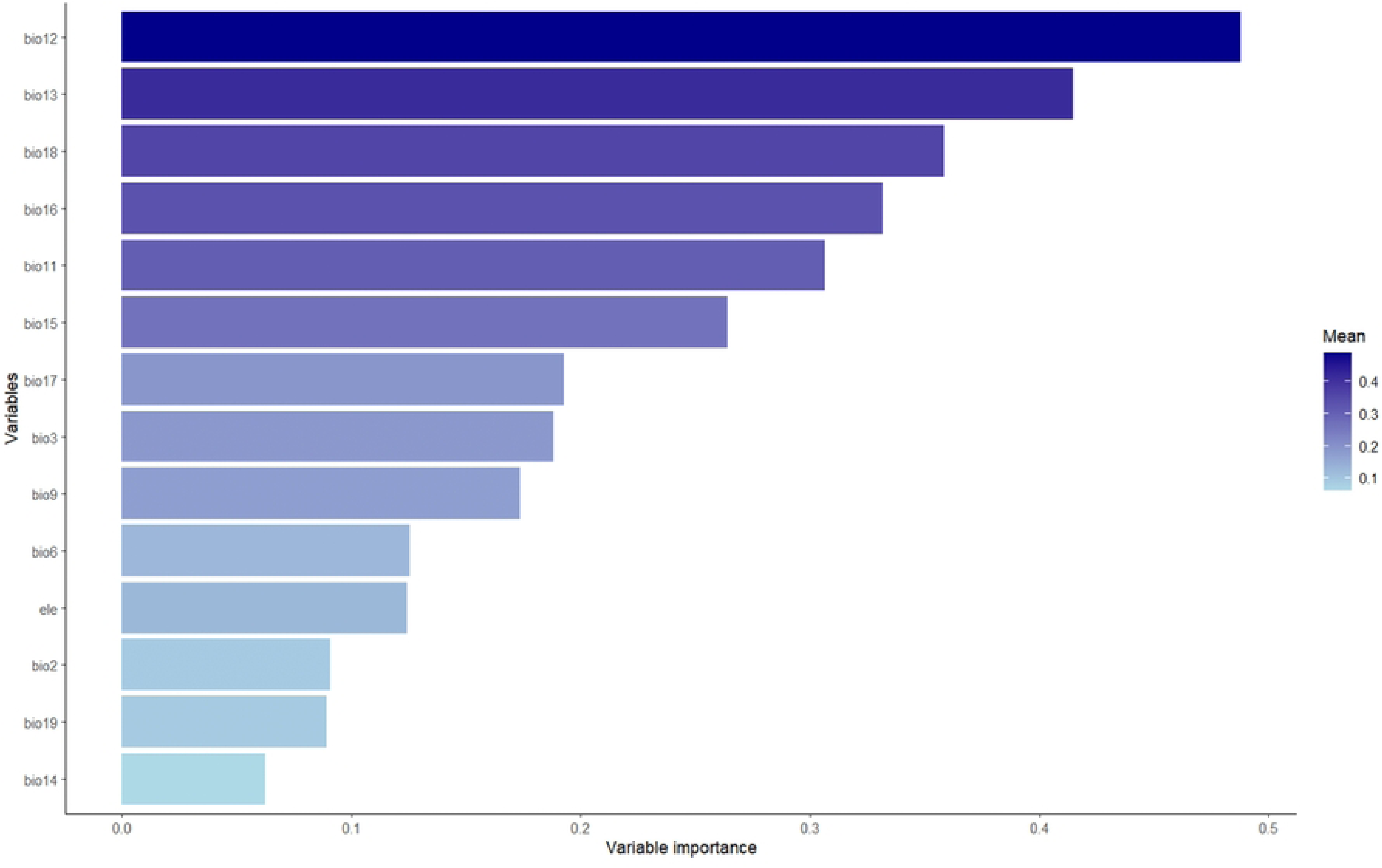
Variable importance plot of all the variables used for modelling.

**Fig 5.**
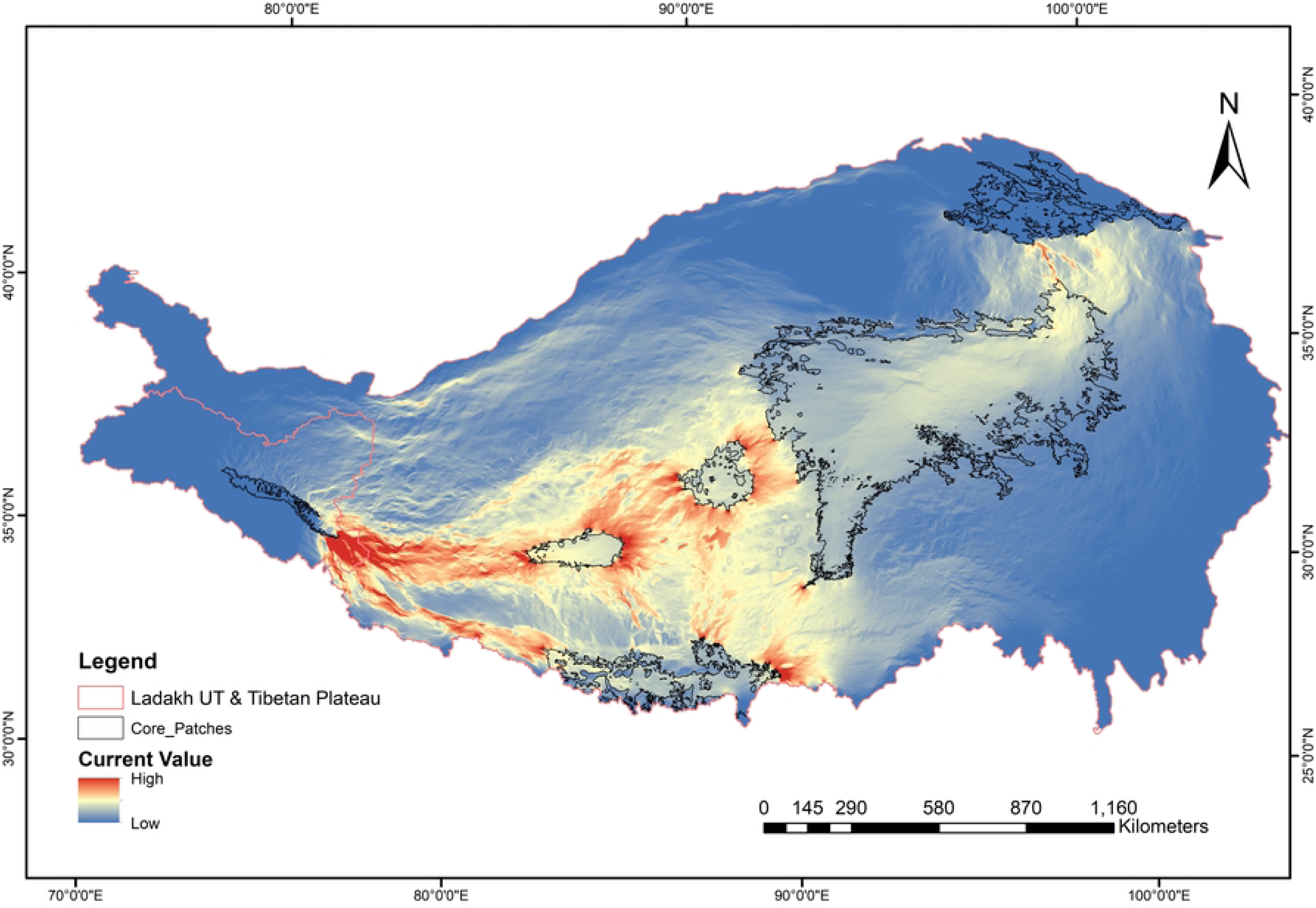
Transboundary Ecological corridors under current climate scenario between Ladakh UT, India and Tibetan plateau

## Discussion

Using an ensemble species distribution modelling framework, we identified suitable habitat for the Tibetan brown bear under current climate scenario across the transboundary landscape encompassing the Tibetan Plateau and the Changthang region of Ladakh, India. Our findings shows that approximately 1011818 km^2^ (21.34%) area combining Ladakh UT, India and Tibetan plateau are currently suitable for Tibetan brown bear. However, out of this current suitable habitat, only 207000 km ^2^ areas come under high suitable areas (Fig 3). Among all predictors, annual precipitation (Bio12) and precipitation of the wettest month (Bio13), Precipitation of warmest quarter (Bio18) emerged as the most influential variables affecting Tibetan brown bear current habitat suitability (Fig 4). Present finding underscores the importance of moisture availability in shaping current distribution of Tibetan brown bears in cold desert ecosystems, where primary productivity and prey availability are strongly constrained by precipitation. Previous studies conducted on Tibetan brown bear highlighted that change precipitation patterns and temperature shift may affect the habitat suitability of the species [26,29,59]. Precipitation and temperature are widely recognized as key bioclimatic factors shaping the geographic distribution and persistence of brown bears across distribution range. Alteration in precipitation not only shift bears distribution to climatically suitable habitats, but also increase the risk of range contraction or fragmentation that may threaten population viability [10]. In high-altitude landscapes, precipitation plays a particularly complex role, while stable annual precipitation can enhance primary productivity and resource availability, however, the higher precipitation during the wettest periods may impose seasonal constraints by deeper snow cover leads to less resource availability and can affects movement costs [88,89]. Increased energetic cost associated with higher movement speeds and longer foraging distances under wetter conditions may further increase metabolic costs, especially during high energy requirement phases [89]. Moreover, poor precipitation during winter can leads to low snow covers can affect denning patterns and post-denning habitat conditions during spring and summer due to change in vegetation structure [25], with potential consequences for reproductive success and cub survival [90]. Significant influence of precipitation variables observed in this study, highlighting how changes in precipitation patterns (annual and seasonal) can shape habitat suitability and connectivity for brown bears in high altitude regions. The studies have shown in high altitude areas are susceptible to warming affects because of optimal temperature and stable precipitation [91]. However, due to climate change, alteration in precipitation and temperature could change the vegetation composition thus affecting habitat distribution [92]. Climate warming is projected to occur at an average global velocity of approximately 0.42 km per year (range: 0.11–0.46 km per year) [93], and species are therefore expected to adjust by shifting their geographic ranges in order find suitable climatic conditions [8,94]. There are various studies which shows impact of future climate change scenario on the habitat fragmentation, distribution and range shift of brown bear [25–29,31–33,59]. Therefore, identifying the ecologically functional corridor is crucial for facilitating movements, dispersal, gene flow and climate induced range shift [95–99]. Findings our connectivity analysis revealed ecological corridors linking suitable habitat patches in the Tibetan Plateau with the recently documented occurrence sites in the Changthang region. Areas of high current flow identified using Circuitscape suggest areas where landscape configuration may facilitate Tibetan brown bear movement, indicating that Changthang is not ecologically isolated but may be functionally connected to broader Tibetan brown bear habitats. These findings are consistent with previous connectivity studies on wide-ranging carnivores, which have shown that circuit models are effective for identifying corridors across heterogeneous landscapes [48,87,97]. For species such as brown bears, which exhibit long-distance dispersal, exploratory movements, and climate driven range shift this approach is especially applicable. (26,28,29,32,59].

Recent record of Tibetan brown bear from Changthang region raises several ecological interpretations. One possibility is that this record may reflects a long-distance dispersal event from source populations on the Tibetan Plateau, facilitated by existing corridors. As evident from previous studies, Brown bears are known to undertake extensive natal dispersal, often exceeding several hundred kilometres, particularly in low-density populations [100,101].

Alternatively, the occurrence may indicate a gradual range expansion or reoccupation of previously suitable but undocumented habitats. Climate change has been widely documented to drive shifts in species geographic distributions, particularly in montane and high-latitude systems [9,11]. In the high-altitude Trans-Himalayan landscape, changes in temperature and precipitation may be altering habitat suitability areas, and enabling bears expansion to new areas where suitable climatic conditions were previously prohibitive [3,26,32]. A third possibility is that Tibetan brown bears may have historically occupied parts of Changthang but remained undocumented due to extremely low population densities. However, the spatial connectivity between suitable habitat patches across transboundary landscape and areas of high current flow provides a confirmation that the Changthang record of Tibetan brown bear existence landscape connectivity rather than a case of isolated dispersal event. Additionally, the evidence from camera trap indicates that Tibetan brown bears in the Changthang region were involved in conflicts with local communities.

We recommend detailed systematic monitoring of the Tibetan brown bear in this region, using robust methods such as camera traps, sign surveys, and non-invasive genetic sampling. A comprehensive monitoring study will provide further insights into its occurrence and enhance our understanding of its range expansion.

## Acknowledgment

The authors sincerely thank the Department of Wildlife Protection, Ladakh, India, for providing the species data and supporting the field surveys. We are grateful to the Director, Zoological Survey of India, for encouragement and institutional support

## Supporting information

S1Table. Environmental predictors used to model habitat suitability and their contribution

S1 Fig. Pearson correlation plot of all the predictors used for modelling (variables with values

S2 Fig a. Response curve of glm model used for modelling

S2 Fig b. Response curve of gam model used for modelling

S2 Fig c. Response curve of RF model used for modelling

